# Epigenetic adaptation prolongs photoreceptor survival during retinal degeneration

**DOI:** 10.1101/774950

**Authors:** Rachayata Dharmat, Sangbae Kim, Hehe Liu, Shangyi Fu, Yumei Li, Rui Chen

## Abstract

Neural degenerative diseases often display a progressive loss of cells as a stretched exponential distribution. The mechanisms underlying the survival of a subset of genetically identical cells in a population beyond what is expected by chance alone remains unknown. To gain mechanistic insights underlying prolonged cellular survival, we used *Spata7* mutant mice as a model and performed single-cell transcriptomic profiling of retinal tissue along the time course of photoreceptor degeneration. Intriguingly, rod cells that survive beyond the initial rapid cell apoptosis phase progressively acquire a distinct transcriptome profile. In these rod cells, expression of photoreceptor-specific phototransduction pathway genes is downregulated while expression of other retinal cell type-specific marker genes is upregulated. These transcriptomic changes are achieved by modulation of the epigenome and changes of the chromatin state at these loci, as indicated by immunofluorescence staining and single-cell ATAC-seq. Consistent with this model, when induction of the repressive epigenetic state is blocked by *in vivo* histone deacetylase inhibition, all photoreceptors in the mutant retina undergo rapid degeneration, strongly curtailing the stretched exponential distribution. Our study reveals an intrinsic mechanism by which neural cells progressively adapt to genetic stress to achieve prolonged survival through epigenomic regulation and chromatin state modulation.

## Introduction

Neurodegenerative disorders of the central nervous system are a group of debilitating diseases characterized by severe physical, cognitive, and visual defects based on the cell type affected. Extensive research over the past several decades has significantly improved our understanding of the disease etiology. Currently, the causative molecular lesion can be identified for approximately 10% of neurodegenerative diseases of the brain and 60% of photoreceptor degenerative diseases (Pang et al., 2017; Richards et al., 2018). A common feature of neurodegenerative diseases is the progressive death of neurons over time where the kinetics of cell death have been described to follow a stretched exponential model (Clarke and Lumsden, 2005; Torriglia et al., 2016). This characteristic and progressive cell loss over a prolonged period of time is observed in both mouse models and human patients (Clarke et al., 2000; Clarke and Lumsden, 2005; Eblimit et al., 2015; Hong et al., 2000). For example, *SPATA7* has been identified as the *LCA3* disease gene, mutations in which lead to early onset photoreceptor degeneration in humans (Wang et al., 2009). Photoreceptor cell degeneration in *Spata7* mutant mice can be separated into two phases: an initial rapid degeneration phase, where 40% of photoreceptor cells die within the first two weeks after eye opening, followed by a slow degenerative phase where about 20% photoreceptor can survive beyond 6 months of age (Eblimit et al., 2015). It is a conundrum how a certain proportion of cells succumb rapidly during the timecourse of degeneration, while other genetically identical cells in the population prolong their survival well beyond what is expected by chance alone until the tail end of the degeneration timecourse. It has been hypothesized that cellular variability in a clonal population can lead to a varied response to intrinsic and extrinsic insults (Galluzzi et al., 2018). However, the validity of this hypothesis and the underlying mechanisms that prolong survival of a subset of cells in a genetically identical population have not been investigated.

Similar to other parts of the central nervous system, the onset and the time course of photoreceptor neuron degeneration is determined by the interplay between the local microenvironment and the underlying genetic mutation. Apoptosis of photoreceptors is controlled by two competing forces: pro-apoptotic factors and neuroprotective signals. In photoreceptors, different mutations often converge on a common set of pathways such as protein mislocalization, protein misfolding, ER stress, and mitochondria malfunction with accumulation of reactive oxygen species (ROS) (Bramall et al., 2010; Doonan et al., 2012; Pacione et al., 2003; Samardzija et al., 2012). While some of these features are pro-death, such as toxic accumulation of mistrafficked Rhodopsin (Nemet et al., 2015; Sung and Tai, 2000), others are pro-survival factors, such as STAT3, which is neuroprotective when upregulated in Müller glia (Jiang et al., 2014). Shifting the balance of these two competing forces could determine whether any given cell follows a path of survival or apoptosis. Therefore, a better understand of the underlying mechanisms that control this balance may result in novel therapeutic avenues for this largely untreatable group of inherited disorders.

Conceptually, prolonged cell survival during neurodegeneration can be explained by at least two models. First is a “selection model,” where the difference in cell death is due to subtle differences in fitness among individual cells in the population. Cells that are more fit can survive longer. This is analogous to a heterogeneous tumor where multiple clones with different mutations exist. The second is an “adaptation model,” where all cells start out the same, but cells that can adapt to stress better will survive longer. One mode of cellular response to the environment is altering chromatin accessibility through modulation of DNA methylation and histone modification profiles, resulting in changes in transcription dynamics. The most prominent example is cancer cells. Cancer cells can develop resistance to chemotherapy by aberrant expression of drug transporters through epigenetic changes (Zeller and Brown, 2010) (Wilting and Dannenberg, 2012). However, unlike the heterogeneity observed in cancer cells, the epigenome profile is relatively uniform and stable within the same neuronal cell type. To distinguish these two “selection” versus “adaptation” hypotheses, we performed a single-cell transcriptomics-based assessment of gene expression at several time points during the course of photoreceptor degeneration in the *Spata7* null mutant mice. Our results suggest that surviving rod cells gradually alter their transcriptome profiles, with downregulated expression of photoreceptor-specific phototransduction genes and upregulated expression of markers of other retinal cell types. Moreover, profiles of a panel of histone modification markers reveals that surviving photoreceptors adopt a repressive epigenomic state. scATAC-seq further confirms that alteration of gene expression likely results from changes in chromatin accessibility. Suppression of this repressive chromatin profile at the onset of degeneration by histone deacetylase (HDAC) inhibition leads to rapid photoreceptor degeneration. Therefore, our results support an adaptation model where neural cells under stress can achieve better survival by activating intrinsic machinery to progressively change gene expression profiles through epigenomic modifications.

## Results

### Single-cell transcriptome profiling reaveals the retinal composition during degeneration

To probe the underlying mechanism that aids prolonged cell survival in a genetically identical population of degenerating neurons, we chose to profile degenerating photoreceptors in the retina as a model system. Specifically, we chose the mouse model of an inherited retinal dystrophy gene, *SPATA7*, the causal gene for *LCA3* (Wang et al., 2009). In *Spata7* null mutant mice, loss of photoreceptors begins as early as postnatal day 15 (P15) with as much as 40% of the photoreceptors lost by P35 (Eblimit et al., 2015). Thereafter, a slower phase of cell death follows with approximately 10-20% photoreceptors surviving beyond 6 months of age (Sup. Fig. 1). Given these two phases of cell degeneration where a subset of cells survive longer than other genetically equivalent cells, we investigated the transcriptome of degenerating photoreceptors at the single-cell level in *Spata7* mutant mice at multiple time points including P30 (1M, peak cell death), P95 (3M, intermediate timeline), and P180 (6M, surviving/ prolonged cell survival) with age-matched wild-type (WT) controls. Suspensions of live, single cells from the retina were captured using the SMARTer ICELL8 single-cell nanogrid system which permits the capture of up to 1,500 live single cells per chip (Fig. 1A). To avoid technical variability, wild-type and *Spata7* mutant retinal cells were captured on the same chip so that cells from each paired-sample were processed with the same methodology.

**Fig 1:**
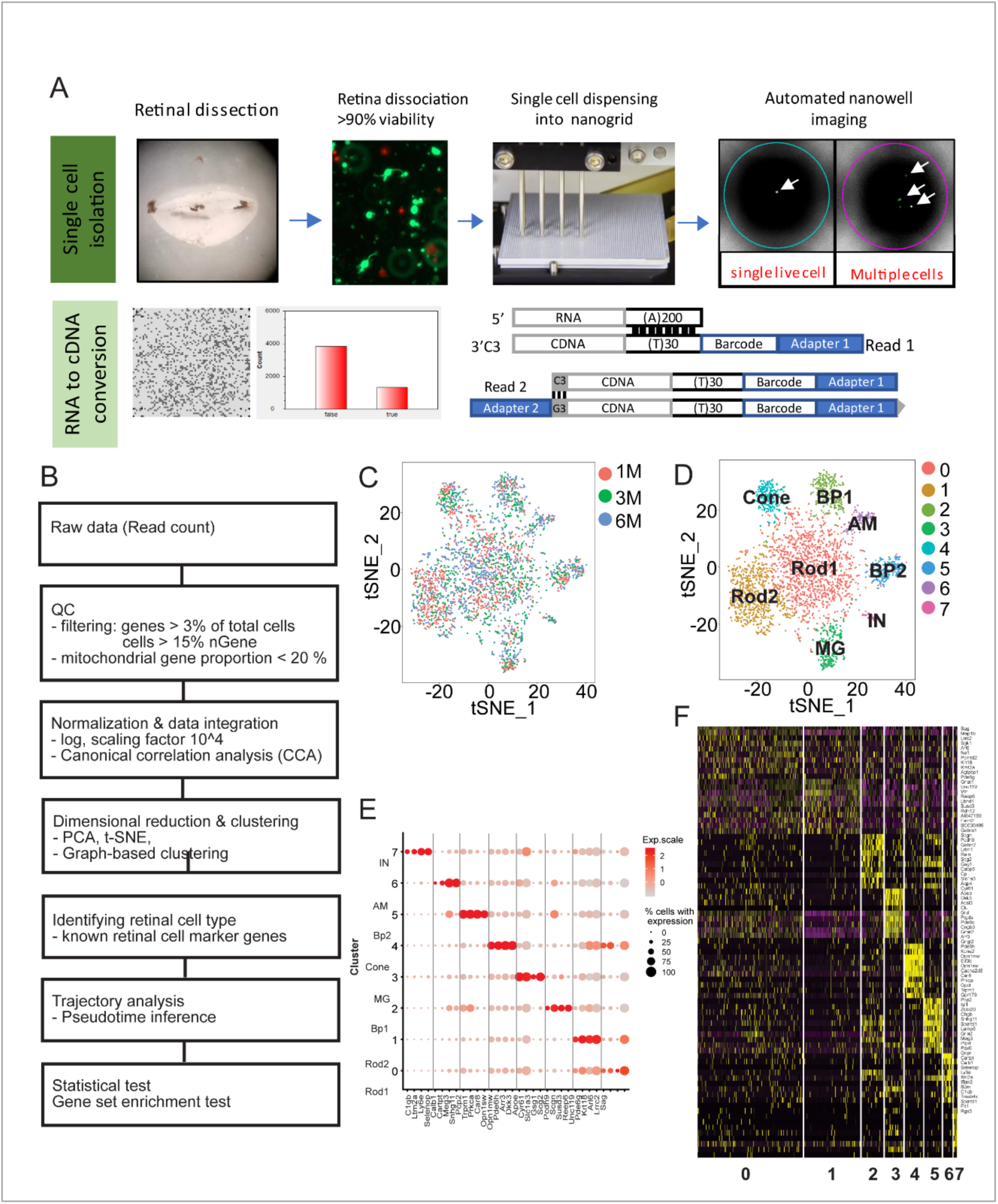
Single cell RNA sequencing of the mouse retina generates 8 distinct cellular clusters. A. Single cell capture and sequencing workflow: Live retinal neurons were captured on an ICELL8 MultiSample NanoDispenser (MSND), and candidate wells selected for downstream RT have a single live cell per well (arrow). Wells containing multiple cells, or debris are discarded from the candidate list. For RNA to cDNA conversion, cell select software generates a map (left panel) for candidate wells (approximately 1500) which are quantified in the right panel which utilizes SMART seq based cDNA conversion (See Methods for more details). B. Workflow of scRNA-seq analysis. C. t-SNE plot of single cells from *Spata7* mutant and wild-type mouse retinas at 1, 3, and 6 months of age. D. Seven clusters from 3,747 cells were identified and assigned to retinal cell types based on known cell-type markers using t-SNE. Each dot indicates a cell and clusters are color-coded. AM, amacrine cell; BP, Bipolar cell; MG, Müller glia; IN, mixed neuron and microglia population. E. Dot plot for top 4 DEGs in each cluster. The color of each dot indicates average scale expression from low in grey to high in red, and its size represents the percentage of cells with expression in the cluster. F. Heatmap of the top 10 differentially up-regulated genes of each cluster compared with others. Expression levels range from high in yellow to low in purple.

After quality control and filtering as described in the Methods section, single-cell transcriptome profiles for 3,743 cells across the three time points for both genotypes was obtained. This dataset included 1,510 cells from 1M, 1,013 cells from 3M, and 1,224 cells from the 6M time points. The scRNA-seq analysis was performed as outlined in the workflow (Fig. 1B). In brief, to generate an integrated dataset, we aligned all data sets using canonical correlation analysis (CCA) in Seurat and performed non-linear dimensionality reduction using t-distributed stochastic neighbor embedding (t-SNE). After aligning the datasets, we confirmed that cells from three different time points were evenly distributed implying lack of batch effects (Fig.1C, Fig2A). Since retinal cells are post-mitotic terminally differentiated cells, the impact of the cell-cycle phase on clustering assignments was negligible, with majority of cells observed in the G2M phase as expected (Sup. Fig 2). To classify cell types during the time course of degeneration, we identified 8 clusters using the graph-based clustering method in the Seurat package and visualized these clusters using t-SNE. The top 3 and top 10 upregulated genes of each cluster were displayed using dot plot and heatmap, respectively (Fig.1D, E, F). Based on the expression of known cell type-specific markers, each cluster was manually assigned to the corresponding retinal cell-type. As expected, all major retinal cell types, including rod, cone, rod bipolar (rodBP), cone bipolar (coneBP), Müller glia (MG), amacrine cells (AM), and a mixed cell population (IN) representing the remaining cell types (horizontal cells, microglia, and endothelial cells) were identified (See Sup. Table 1 for markers used and references). Clusters 0 and 1 were identified as Rods using *Sag*, *Pde6a*, *Nr2e3, Pde6g*, *Gnat1*, and *Reep6* as markers. We assigned other clusters using known markers as followings: *Scgn, Grm6, Grik1*, and *Vax1* for cluster 2 as coneBP; *Apoe*, *Dkk3*, and *Glul* for cluster 3 as MG; *Pde6c*, *Arr3*, and *Opn1mw* for cluster 4 as cone cells; *Prkca, Bhlhe23*, and *Car8* for cluster 5 as rodBP; *Calb1*, *Cartpt*, and *Pax6* for cluster 6 as AM; and cluster 7 as a mixed population of horizontal cells (identified by *Lhx1, Onecut1, Onecut2*, and *Calb1*), microglia (identified by *C1qa, C1qb, C1qc, Hexb*, and *Ctss*), and endothelial cells (*Cldn5, Igfbp7*, and *Col4a1*). Since the rod photoreceptor cell is the major cell type in the retina, it is not surprising that about 65% of the cells identified in our dataset were indeed rod (Fig. 2B). Interestingly, although a single type of rod cell is present in the wild-type mouse retina (Macosko et al., 2015; Shekhar et al., 2016), two distinct clusters of rods were observed in our dataset (Fig. 1D), suggesting transcriptional profile heterogeneity in rods. Given that *Spata7* is required primarily for rod photoreceptor cell survival, we further investigated if the transcription heterogeneity was related to genotype (Fig. 2A-C). Indeed, the two rod clusters strongly correlated with genotype where the first rod cluster (Rod1) is comprised primarily of *Spata7* mutant cells (95%) while the second rod cluster (Rod2) is mostly wild-type (85%). Thus, unlike other cell types, distinct expression profiles are evident in rods isolated from *Spata7* mutants versus wild-type controls.

**Fig 2:**
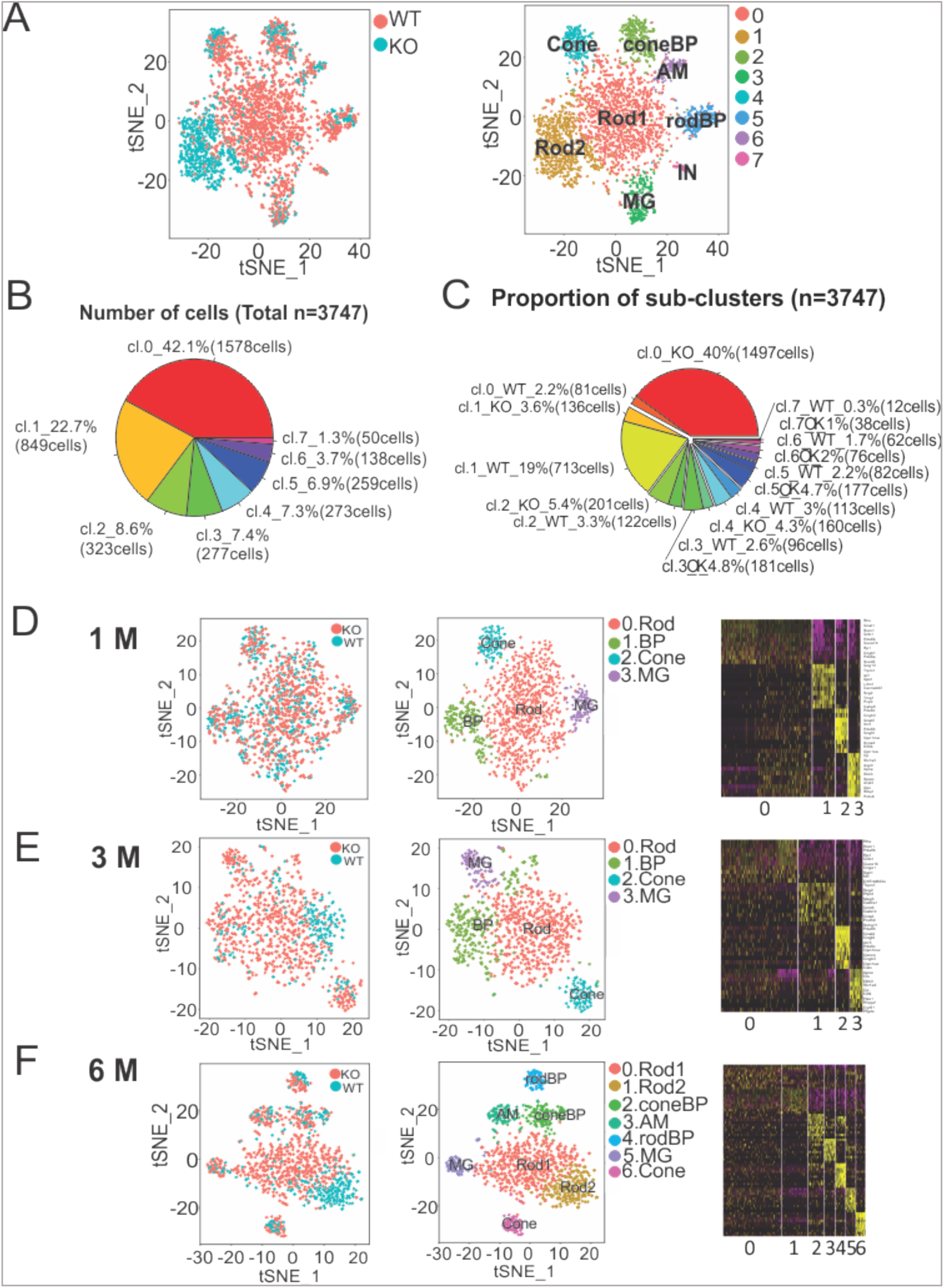
*Spata7* mutant rods display an increasingly distinct transcriptional profile from wild-type. A. t-SNE plots of the integrated dataset were visualized by genotype and cluster identity displaying distinct clusters of wild-type and degenerating rod cells. B. Proportions of cells were calculated for 7 clusters. C. The proportion of sub-groups assigned by genetic condition (WT or KO) were calculated. D, E, and F. t-SNE plot of single cells from *Spata7* mutant and wild-type mouse retinas at 1, 3, and 6 months, respectively, using the Seurat package. Left panels show t-SNE plot by genotype. Middle panels represent t-SNE plots to identify 4 (1 and 3 month) and 7 (2 month) retinal cell-type clusters. Each dot indicates a cell and clusters are color-coded. AM, amacrine cell; BP, Bipolar cell; MG, Müller glia; IN, intermediate neural cell. Right panel shows the heatmap of the top 10 differentially up-regulated genes of each cluster compared with others at 1 month (D), 3 months (E), and 6 months (F). Gene expression levels range from high in yellow to low in purple.

### Mutant Rods display an increasingly distinct transcriptome profile from wild-type

Differences in transcriptome profiles between surviving *Spata7* mutant and wild-type control rod photoreceptors could either exist from the onset of degeneration or accumulate progressively over time. To distinguish these two scenarios, we analyzed the expression profiles of wild-type and *Spata7* mutant rods across the degeneration timeline (Fig 2. D-E). At 1 month, instead of two rod clusters, cells from the two genotypes mixed evenly and resulted in one rod cluster, indicating that at the peak of cell death the mutant and wild-type rods have similar profiles (Fig. 2D). In contrast, at 3 months, although the *Spata7* mutant and wild-type rods cluster together, segregation of these two genotypes within the cluster is observed (Fig. 2E). The segregation is more evident at 6 months and results in two distinct rod clusters, where the Rod1 cluster is comprised primarily of surviving *Spata7* mutant rods while the wild-type rods are enriched in the Rod2 cluster (Fig. 2F). Therefore, our time series data suggest that although similar at the beginning, the rod cell expression profiles of wild-type and *Spata7* mutant rods become progressively distinct over time.

### Mutant rods significantly deviate from the wild-type photoreceptor-specific expression profile

To investigate the differences between wild-type and *Spata7* mutant rods, differentially expressed genes (DEGs) were identified by the Wilcoxon rank sum test for all three time points. A total of 460 DEGs were identified among the 6 datasets, including 283 genes at 1 month, 97 genes at 3 months, and 236 genes at 6 months of age (Sup. Fig. 3). Among all the DEGs (adj value <0.05), 41 genes were common across all datasets (Fig. 3A and B). As expected, several stress and repair response related genes were consistently upregulated across multiple time points. For example, expression of genes acting in the VEGF signaling pathway (*Kdr*, *Cd63*, and *Tcf4*), G-protein receptor signaling (*Apoe*, *Gnb3*, *Gng13*, and *Opn1sw*), axodendritic transport (*Kif1a*, *App*, *Nefl*, and *Kif5a*) and glial proliferation signaling (*Glul*, *Dbi*, and Vim) are induced in *Spata7* mutants (Clifford et al., 2008; Gogat et al., 2004; Huang et al., 2003; Li et al., 2014; Ramakrishnan et al., 2015; Roesch et al., 2008). Moreover, many genes that are specific markers for non-rod retinal cell types are upregulated across the three time points. For example, misexpression of markers of bipolar cells (GNB3 and PCP2), Müller glia (*Apoe, Clu, Rlbp1*, and *Glul*), cone cells *(Pde6h*), and genes associated with active proliferation (*Dkk3*) are observed in *Spata7* mutant rod cells (Fig 3B, 3F), thus displaying a non-lineage specific expression pattern in surviving rods (Roesch et al., 2008; Shekhar et al., 2016). To confirm these findings, antibody staining for GNB3, which is normally expressed in bipolar cells, was performed (Huang et al., 2003). As shown in Figure 3H, increased and aberrant localization of GNB3 is observed in the outer nuclear layer of *Spata7* mutants compared to wild-type.

**Fig 3:**
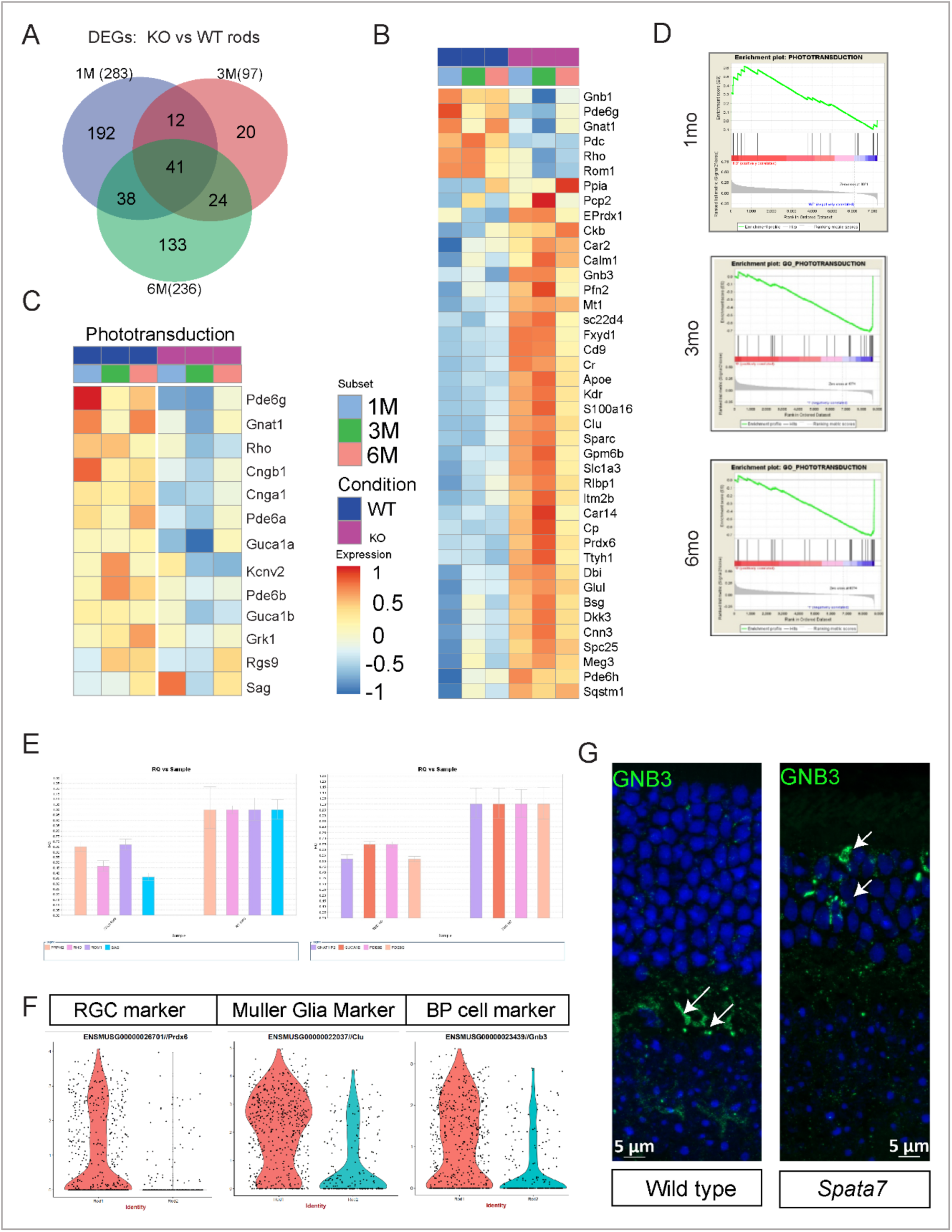
Phototransduction pathway genes are downregulated in *Spata7* mutant rod cells. A. DEGs between subclRTusters were identified using the differentialGeneTest function of Monocle2 with FDR < 0.01. A Venn diagram shows the overlap between the two comparisons. B. Heatmap displaying the average expression of sub-groups by condition and subset for 41 common DEGs across the aggregate dataset. Expression values are median centered after log2 transformation and scaled to display Z scores. C. Average expression of genes of sub-groups by condition and subset in phototransduction was visualized with heatmap. Expression values are median centered after log2 transformation. D. Gene Set Enrichment Analysis (GSEA) plot of age-matched dataset comparison displayed an increasingly significant downregulation of the phototransduction pathway genes with increase in age. E. qRT-PCR analysis of gene expression in *Spata7* mutants and wild-type at 6 mo of age normalized against GAPDH expression (N=3). F. Violin plots for expression of markers of non-photoreceptor cell markers in rod cells in *Spata7* mutants and wild-type at 6 mo. G. Immunofluorescence localization displays aberrant/increased localization of GNB3 in photoreceptor outer nuclear layer in *Spata7* mutants compared to wild-type at 6 mo.

Among the downregulated DEGs, the most significantly enriched set of pathways are photoreceptor cell-specific pathways such as visual perception and phototransduction (Sup. Fig 3). Indeed, the expression level of many genes involved in rod photoreceptor cell phototransduction are significantly reduced in *Spata7* mutants across all the three time points (Fig 3B). We thus specifically assessed the known phototransduction genes beyond those discovered among the shared 41 genes across the integrated dataset and observed a significant downregulation (Fig 3C). Furthermore, just as the overall transcriptome profiles of *Spata7* mutant and wild-type rods become increasingly distinct over time, the differential expression of phototransduction pathway genes also become increasingly significant over the timeline of degeneration, as shown in Figure 3D, using the Gene Set Enrichment Analysis (GSEA) algorithm. By 6 months, most of the known rod phototransduction genes are significantly downregulated, including *Rho, Gnat1, Gnb1, Cngb1, Grk1, Gngt1, Guca1b, Pde6g*, and *Rom1* (Fig 3C) (Larhammar et al., 2009). This observation was further verified by qRT-PCR using RNA prepared from retinas of 6 month old *Spata7* mutant mice, where a 2-fold reduction was observed in all phototransduction genes tested (Fig 3E). Downregulation of phototransduction gene expression level can indeed have neuroprotective effects and promote cell survival. We have previously shown that *Spata7* is required to maintain the proper structure and function of the connecting cilium, an organelle which plays a critical role in protein transport between the inner segment and the outer segment of rods (Dharmat et al., 2018). In *Spata7* mutants, rhodopsin and other highly expressed phototransduction proteins accumulate in the inner segment of rods, resulting in ER-stress and cellular toxicity. Downregulation of phototransduction genes, such as *Rho*, can prolong cellular survival (Eblimit et al., 2015). Therefore, the downregulation of transcription across several phototransduction genes is likely to offer a protective effect for *Spata7* mutant rods, and this mechanism is likely to at least partially account for the prolonged cell survival phenotype we observe.

To delineate the transcriptional dynamics of prolonged cell survival, we generated pseudo-time trajectories for 1 month, 3 month, and 6 month rod cells using wild-type (794) and *Spata7* mutant (1,633) rods extracted from the integrated analysis, encompassing 7 different pseudotime stages that branch into four ends (Sup. Fig 4A). About 89% wild-type and 11% of *Spata7* mutant rods rods at 1 month occupy the start of the trajectory (C1) (Sup. Fig 4B-C). The majority of *Spata7* mutant rods at 1 month are specifically enriched along branches C5, C6 and C7 with 18%, 37%, and 31%, respectively. Furthermore, we found a dramatic change in cell proportions among clusters along the timecourse of degeneration (Sup. Fig. 4B-C). In the wild-type, the cell proportion of cluster 1 sharply decreases over time to 6.1%, while that of cluster 3 increases from 5.9% to 72.4%, implying that cells in cluster 1 may represent an early stage while cells in cluster 3 may be a late stage along the degeneration or aging process. Surprisingly, in *Spata7* mutant rods, the proportion of cells in cluster 7 increases dramatically over time, from 31.2% to 88.9%, suggesting that rods with the transcriptome dynamics of cluster 7 are enriched along the timecourse of degeneration (Sup. Fig. 4B-C). This transition of the *Spata7* mutant rods can be a result of modulation of the transcriptome along the timeline of degeneration and as a result transition from a C5/C6-like state to a C7-like state, further indicating towards cellular adaptation.

### Epigenetic modulation changes the cellular transcriptome

We sought to investigate the molecular mechanism for upregulation of non-photoreceptor specific genes and downregulation of photoreceptor-cell specific genes in surviving rod cells. We first tested if the transcriptome profile change could be explained by alterations in the levels of transcription factors that mediate cell type-specific gene expression. A set of transcription factors that play a key role in regulating the development of rods and other retinal cell types were assessed in our dataset. The expression levels of these known transcription regulators, such as *Otx2, Nr2e3, Nrl, Crx*, were largely unchanged or did not show a consistent change across the timeline of degeneration (Sup. Fig. 5) (Brzezinski and Reh, 2015). For example, the expression level of key transcription factors that specify rod cell identity, such as *Otx2*, *Crx*, and *Nr2e3*, remain unchanged across all three time points. Although *Nrl* is downregulated at 1 month, it’s expression in surviving *Spata7* mutant rods was at a similar level to wild-type at 3 months and 6 months. Similarly, expression levels of transcription regulators that are specific to other retinal cell types remain unchanged with a few exceptions (Sup. Fig. 5). For example, *Prox1*, which is the only transcription factor significantly upregulated at both 3 and 6 months, plays an important role in horizontal cell genesis (Dyer et al., 2003). Therefore, while changes in the expression of transcription factors may partially explain the upregulation of other retinal cell type-specific genes in *Spata7* mutant rod cells, it does not explain the downregulation of phototransduction pathway genes.

Given the broad and specific misregulation of cell type-specific pathways, it is conceivable that this is achieved through epigenomic modulation. To test this hypothesis, we profiled a panel of histone modification marks, including active marks, such as pan H4-Ac, H3K27-Ac, H3K9-Ac, and H3K4-Me3, and repressive marks, such as H3K9-Me3 and H3K27-Me3, in surviving rods at the 6 month time point (Zhang et al., 2015). In contrast to the conventional nuclear architecture found in nearly all vertebrate cells, where the heterochromatin is localized at the nuclear periphery and euchromatin is in the nuclear interior, rods cells have a unique inverted nuclear architecture (Solovei et al., 2009). Specifically, rods have a dense heterochromatin region that localizes to the center of the nucleas, whereas euchromatin, where nascent transcripts and splicing machinery are enriched, localizes to the nuclear periphery (Fig. 4A). This allows ready demarcation of the transcriptionally active versus repressed regions of photoreceptors. Normally, active histone marks are detected at the nuclear periphery of rods (Fig. 4B). In contrast, surviving rods in *Spata7* mutants display an overall downregulation of both pan H4-Ac and H3K27-Ac marks. In addition H3K9Ac, which marks promoter regions, is highly upregulated in the heterochromatic region of *Spata7* mutant rods. Consistent with our data, upregulation of H3K9Ac was previously reported to be concomitant with progenitor-like status in photoreceptors and block in Rho expression (Ferreira et al., 2017). Similarly, although repressive marks (H3K9-Me3, and H3K27-Me3) are not detected in wild-type rods, a significant upregulation at the nuclear periphery is observed in *Spata7* mutants (Fig 4B). Therefore, surviving rods appear to have an overall repressive chromatin signature with upregulation of repressive marks and downregulation of active marks within the euchromatin at the nuclear periphery. Given that the nuclear periphery (euchromatin) is the site of active transcription of phototransduction genes such as *Rho* and other cell-specific genes, it is possible that modulation of the cellular transcriptome occurs via epigenetic changes (Solovei et al., 2009).

**Fig 4:**
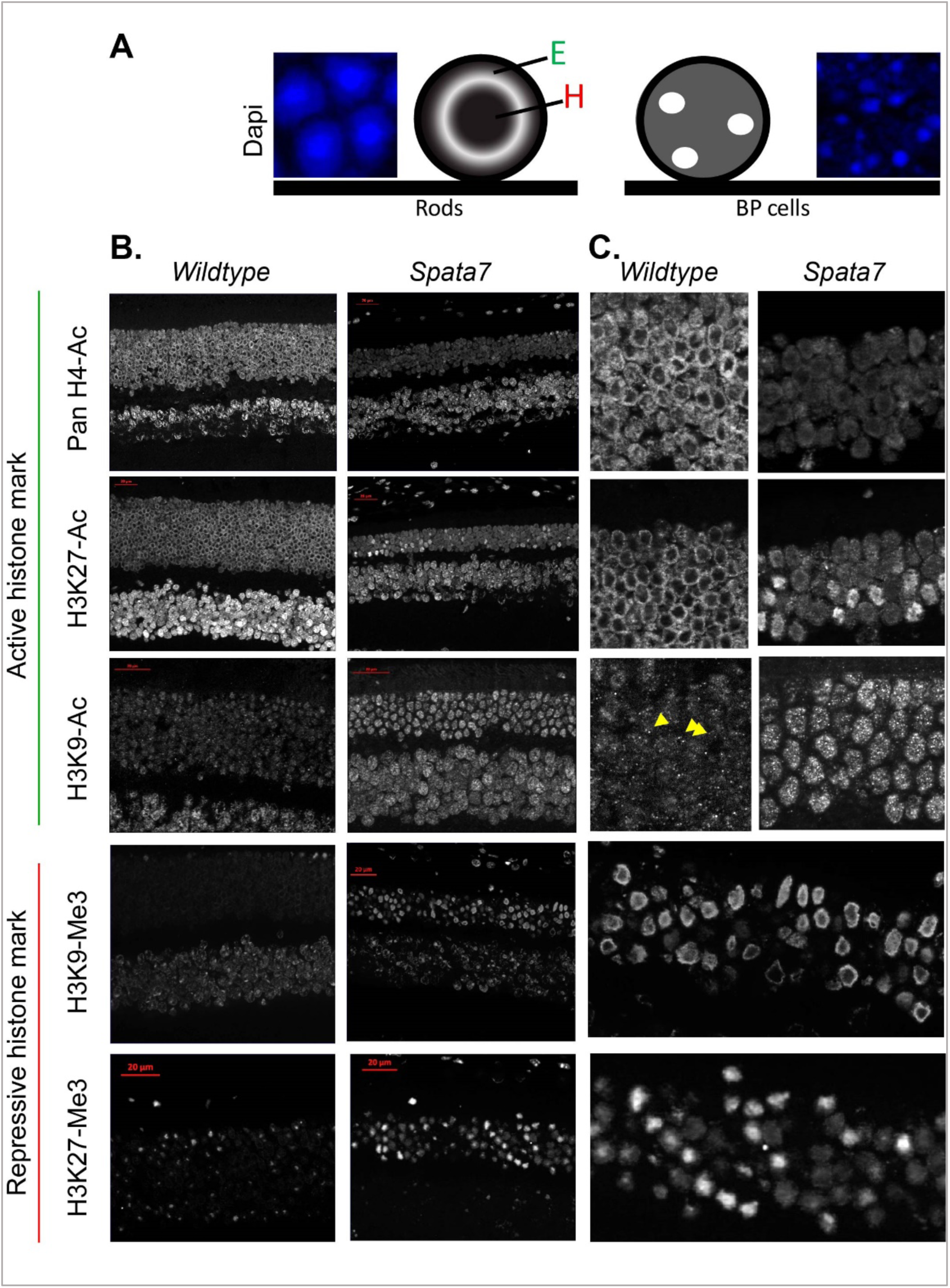
*Spata7* mutant rods display a repressed chromatin profile. A. Inverted nuclear architecture is observed in rod cells with dense central heterochromatin and peripheral euchromatin in comparison to the conventional nuclear architecture found in nearly all vertebrate cells. B. Immunofluorescence localization of active, transcribing and repressed histone marks to heterochromatin and euchromatin displays a decrease in active histone marks and an increase in repressive marks in the rod euchromatin. Scale bar: 20 µm.

### Reduced chromatin accessibility at repressed loci is evident in surviving rods

In *Spata7* mutant retinas, surviving rods exhibit an altered transcriptome with a reduction in the expression of phototransduction genes. Globally, surviving rods also exhibit a repressed chromatin state; however, it remains unclear whether the observed global alteration in chromatin is also affecting DNA accessibility specifically at phototransduction gene loci. Thus, we used single-cell ATAC-seq (scATAC-seq) to examine chromatin accessibility at the target upregulated and downregulated loci in *Spata7* mutant rods at 6 months of age. In our scATAC-seq data, we identified 4 major clusters using the K-means method. To assign retinal cell types, known marker genes were used to attribute clusters to rods (12,838 cells), BPs (3,878 cells), MGs (1,571 cells), and cones (1,263 cells) (Fig 5A, Supp. Fig. 6) (Sup. Table-3). Similar to our scRNA-seq data, we identified two sub-clusters specifically in rod cells that correlate with the genotype of the cell (WT:10286 and KO:2552) suggesting that chromatin accessibility was altered in mutant rod cells compared to wild-type (Fig 5A). In addition, chromatin accessibility at the promoter region of genes in the phototransduction pathway is significantly reduced in *Spata7* mutant rods as observed by GSEA (Nominal p-value < 0.0001). We also observed a reduction in the accessibility of individual candidate genes of the phototransduction pathway, such as *Rho* and *Sag*, in *Spata7* mutant rods; however, upregulated genes such as *Clu* displayed similar levels of accessibility (Fig 5B-C, Sup. Table-4) in *Spata7* mutants. These data further support our hypothesis that epigenomic changes observed in *Spata7* mutant photoreceptors impact the accessibility of candidate genes which, in turn, likely results in changes in the expression levels of these genes.

**Fig 5:**
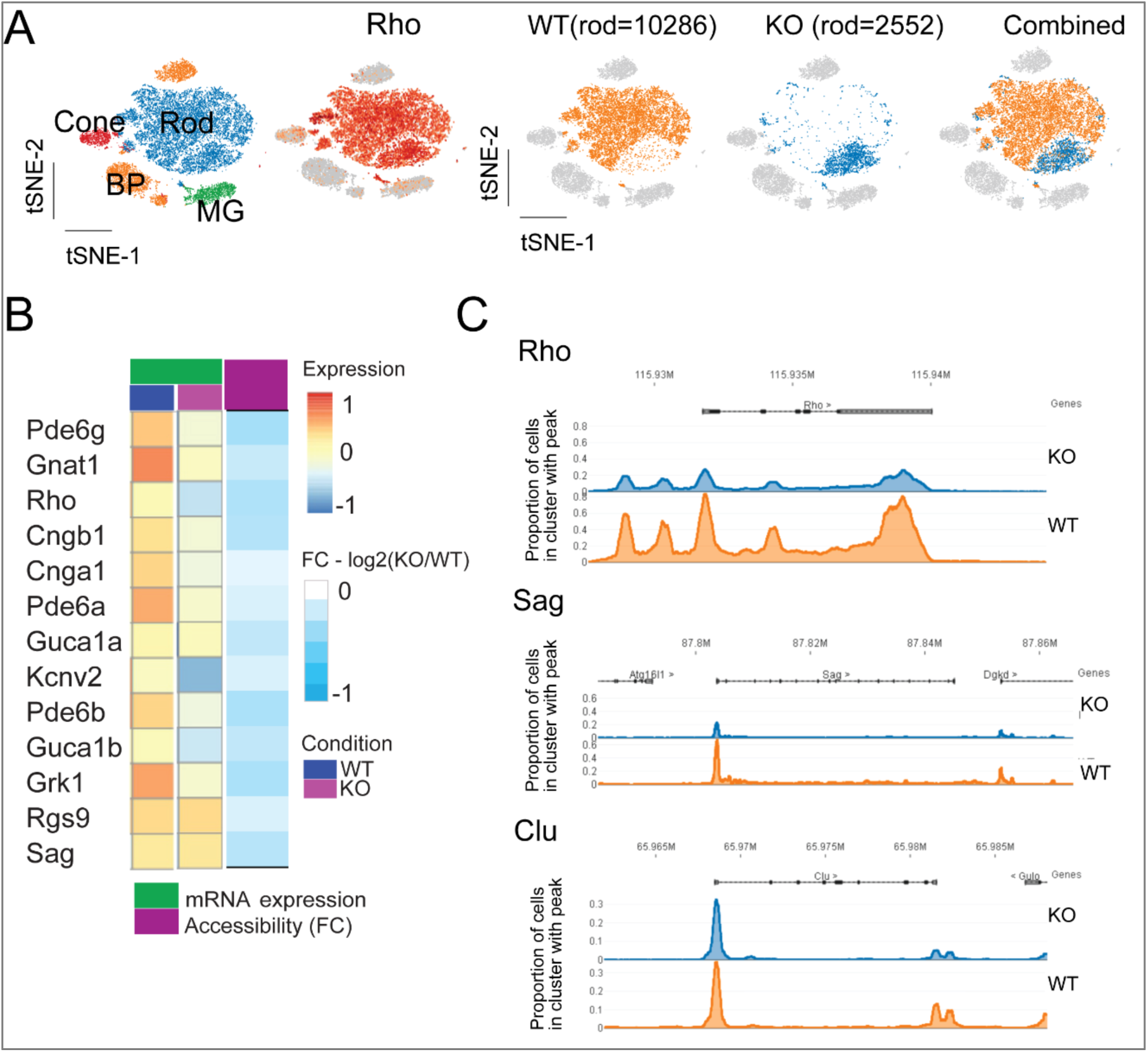
Phototransduction genes have reduced DNA accessibility. A. Single cell ATAC-seq profile: tSNE visualization of rod cluster in 10xscATAC-seq data from WT(n=12985) and *Spata7* KO (n=2552) retinal cells. WT rods in orange and KO rods in blue. B. Chromatin accessibility levels at the promoter regions of genes associated with the phototransduction pathway. C. Average chromatin accessibility levels in the promoter regions of genes related to the phototransduction pathway. Chromatin accessibility of phototransduction pathway associated genes such as *Rho* and *Sag* was lower in *Spata7* mutants in comparison to wild-type. Accessibility of *Clu* however remained unchanged.

### Histone deacetylase inhibition prevents prolonged cell survival in degenerating rods

Our data show a correlation between a neuroprotective signature of downregulated phototransduction pathway genes with a concordant repressed chromatin state in surviving rod cells. It is unclear, however, whether this repressed chromatin phenotype is a secondary event observed in degenerating cells or a primary mediator of prolonged cell survival. To distinguish these two scenarios, we sought to directly prevent a cell from switching to this critical repressed chromatin state by histone deacetylase (HDAC) inhibition. Intravitreal injection of the class I and II HDAC inhibitor trichostatin-A (TSA) preceding the onset of degeneration (P15), followed by a second dose at P21, was performed and the retinal degeneration phenotype was assessed at P30 (Fig 6A) (Kim and Bae, 2011). Both TSA and sham injected contralateral retinas displayed no cell death in the wild-type retina, suggesting little or no cellular toxicity at the dosage of TSA used (660 nM intravitreal concentration). In contrast, accelerated photoreceptor degeneration is observed in TSA-injected *Spata7* mutants (N=4), which retaining only a single row of surviving photoreceptor cells in the outer nuclear layer at P30, compared to the 40-50% outer nuclear layer in contralateral sham injected controls (N=4) (Fig 6B). These data indicate that inhibition of HDAC prevents cellular survival beyond the initial rapid cell death phase. Thus, modulation of the epigenetic profile during retinal degeneration allows cells to adapt and prolong their survival.

**Fig 6:**
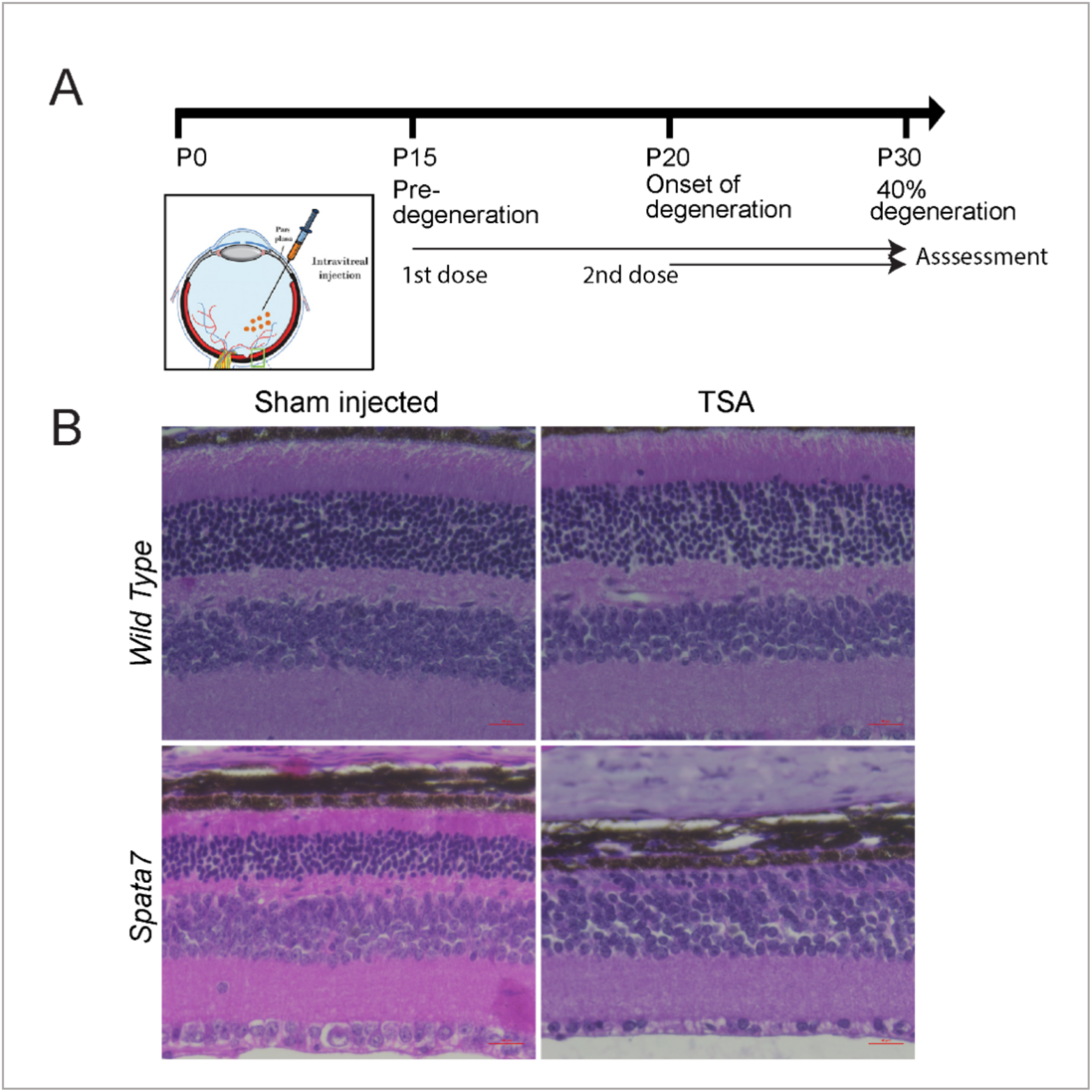
HDAC-inhibition prevents prolonged cell survival of *Spata7* mutant rods. A. Experimental design for intraocular injection of TSA and assessment timeline. B. Hematoxylin and eosin (H&E) staining of paraffin-embedded retinal sections from wild-type (top panels) and *Spata7* mutants where one eye was treated with TSA (660 nM) (N=4/genotype) and the contralateral control eye was treated with sham (50% glycerol). Scale bar: 20 µm.

## Discussion

Although the underlying etiologies of inherited neurodegenerative diseases are becoming more evident over the past decade, variation in cell death kinetics within an affected cell population has not received much attention. Specifically, it remains unclear how a subset of affected cells, despite having an identical genotype and environment, prolong survival beyond that expected by chance alone. This is the first study seeking an answer to this dilemma by directly investigating the heterogeneous cellular and molecular changes at single-cell resolution along the timecourse of degeneration. Single cell RNA sequencing was performed to profile the retinal transcriptome in a mice harboring a null mutation in *Spata7*, an inherited retinal dystrophy gene, across multiple time points along the degeneration timeline. We discovered that *Spata7* mutant rods display an increasingly distinct transcriptomic profile compared to wild-type cells over time. Surviving rod cells exhibit a distinct transcriptomic profile, with downregulated phototransduction pathway genes and upregulated marker genes of several other retinal cell types, including cones, bipolar cells, RGCs, and Müller glial cells. The changes we observed in the transcriptome are likely due to modulation of the epigenome where the euchromatin adopts an overall repressed state. Indeed, a reduction of the accessibility of chromatin around the repressed genes was observed using scATAC-Seq, supporting a direct impact of the chromatin state on modulation of gene expression. Finally, suppression of these epigenetic alterations by histone deacetylase inhibition results in rapid cell degeneration, indicating that epigenetic modification is an intrinsic mechanism the cell adopts to prolong its survival.

An surprising finding of our study is that cells possess intrinsic mechanisms to achieve long-term survival under stress. In *Spata7* mutants, degenerating rods downregulate a large panel of phototransduction pathway genes, including *Rho*, to promote survival. Moreover, the mechanism by which these cellular changes are modulated involves alteration in the cellular chromatin profile, a mechanism that is often observed in cancer cells adapting to gain chemoresistance. To our knowledge, non-cancerous cells, including neurons, have not been reported to employ this mechanism to prolong their cell survival during cellular stress. Such a mechanism is not likely to be unique to the *Spata7* mouse model used in this study as changes in chromatin accessibility have been recently reported as a hallmark of the onset and progression of age-related macular degeneration (AMD), a retinal degenerative disease which has an entirely different genetic underpinning (Wang et al., 2018). In fact, a global and progressive decrease in chromatin accessibility also occurs in retinas of AMD patients (Wang et al., 2018). Indeed, overexpression of HDAC11 in iPSC-derived retinal pigment epithelial cells recapitulates the loss of chromatin accessibility phenotype observed in AMD patients in the study conducted by Wang *et al*. Given the crucial and universal role of epigenetic modifications in gene regulation, we further propose that this intrinsic mechanism utilized by a cell to cope with stress is unlikely to be restricted to the retina. Therefore, further investigation of the role of epigenetic modulation during degeneration is likely to provide important insights and may form the basis for developing novel therapeutic approaches to slow down the progression of an array of neurodegenerative diseases.

Given that global modulation of epigenetic signatures and accessibility of chromatin aids in prolonging cellular survival, observations in our study might have implications for future optimization of degenerative disease therapies. One theory devised to explain stretched exponential cell death kinetics and cellular variability across time is the “mutant steady state” (MSS) (Clarke et al., 2000; Clarke and Lumsden, 2005). According to this theory, mutant neurons are viable but are at an increased risk of initiating the cell death pathway compared to normal neurons. This increased risk factor stems from an increase in the abundance of pro-death molecules which can drive individual neurons past a threshold beyond which the cell ceases to survive. In our study, these pro-death molecules are the set of highly expressed phototransduction proteins. We observe that photoreceptors prolong survival by altering their epigenome and downregulating these pro-death molecules in an effort to adapt and stay below the critical death threshold. Although this adaptation allows cellular survival via global epigenetic modulation, it also impacts the expression of other genes in the cell, thus altering the cellular transcriptome. Moreover, this model also suggests that therapeutic interventions geared toward restoring cellular function may be less effective since surviving cells, although rod-like in nature, display a significantly altered chromatin state. In such cases, therapeutic intervention, such as gene augmentation therapy at a later stage of the disease, may be unsuccessful unless combined with a chromatin modification strategy to restore the epigenetic architecture of surviving rods.

In summary, our study reveals that chromatin accessibility and epigenetic modification is an intrinsic mechanism that allows cellular adaption to stress, thereby promoting long term survival of genetically identical cells. During the initial rapid phase of degeneration, cell apoptosis follows an exponential decay kinetics. As shown in Figure 7, we propose a model where cells that survive this first rapid degeneration phase can progressively adopt an altered state by modifying chromatin accessibility and repressing cell type-specific genes, resulting in a second, slower degeneration phase. Conclusively, reversal of this repressed state prevents prolonged cell survival in the degenerating retina. It is plausible that since cell type-specific genes in a terminally differentiated cell, such as the genes in the phototransduction pathway in the case of rod photoreceptors, are typically turned on later during the development, these genes are more readily subjected to epigenetic regulation. Therefore, modulation of the epigenome can potentially serve as a novel therapeutic approach to reduce the rate of neurodegeneration.

**Fig 7:**
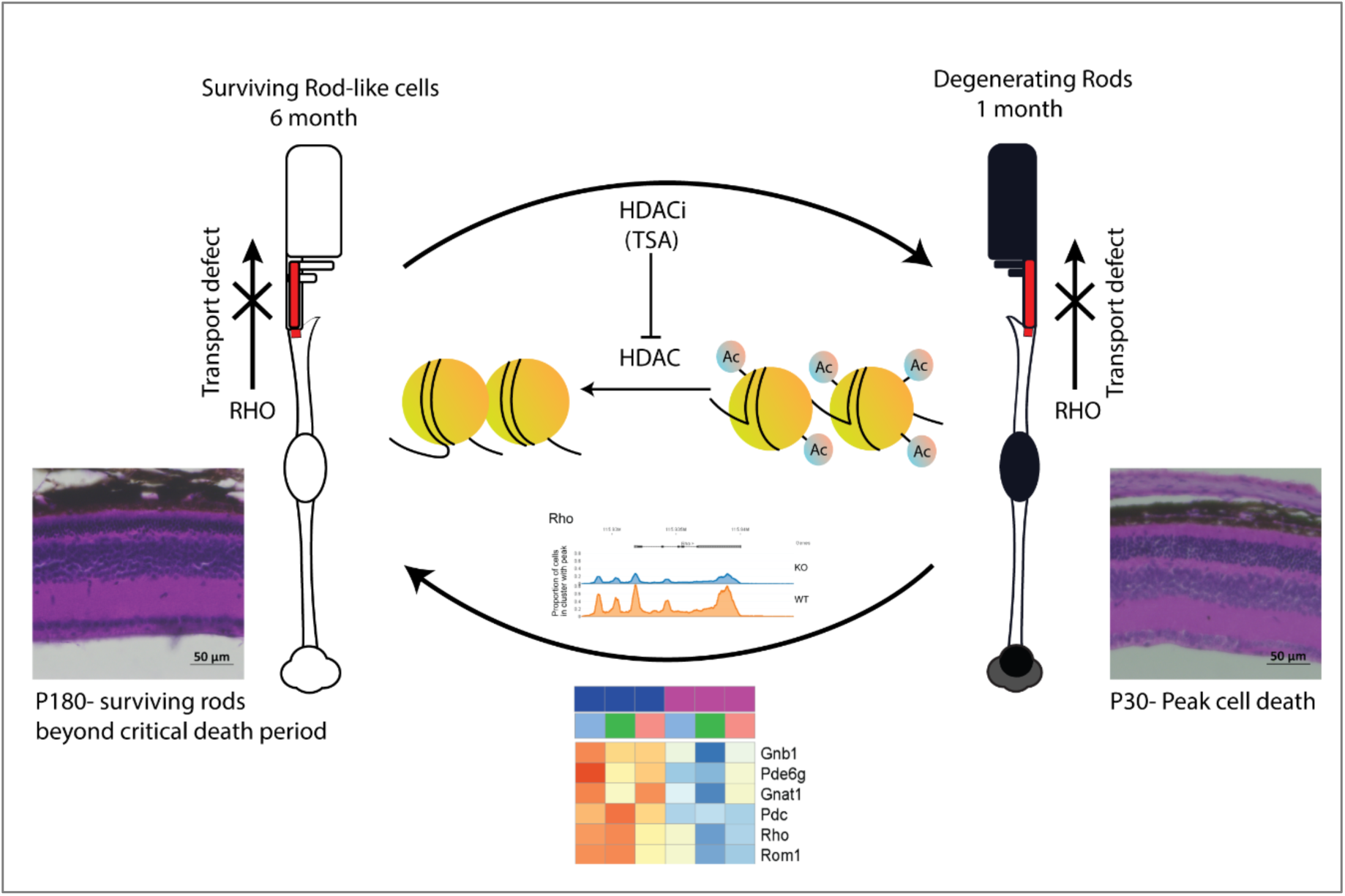
Proposed model for the mechanism of prolonged cell survival in a degenerating retina. Loss of *Spata7* results in a defective connecting cilium structure and this prevents normal transport of proteins into the outer segment of photoreceptor cells, the site of phototransduction, resulting in toxic protein accumulation and cell death. Consequently, the retina undergoes a rapid phase of cell death by P30. We propose a model where a subset of cells can survive beyond the rapid degeneration phase by adopting a repressed state (P180). In this state, expression of rod cell-specific phototransduction genes is reduced by modulation of chromatin accessibility and histone modifications. Reversal of this repressed state in turn prevents prolonged cell survival.

## Materials and Methods

### Animal breeding and mouse models

All animals were handled in accordance with the policies on the treatment of laboratory animals by the Institutional Animal Care and Use Committee of Baylor College of Medicine. Mice were housed on a standard diet and in a 12 hr light to 12 hr dark cycle. For all experiments, male *Spata7* (Eblimit et al., 2015) mutant mice were compared with age-matched male C57BL6/J mice, to avoid sex as a biological variable. All efforts were made to minimize the number of animals used and their suffering.

### Preparation of live single-cell suspensions from the retina

Retinae were harvested from age-matched male C57BL6/J and *Spata7* mutant mice in parallel, while taking precautions to keep the retinal cup intact. Each retina was dissociated using papain-based enzymatic digestion as described previously (Siegert et al., 2012) with slight modifications. Briefly, 45 U of activated papain solution (Worthington, Cat. #LS003126) with 1.2 mg L-cysteine (Sigma) and 1200 U of DNase I (Affymetrix) in 5 ml of HBSS buffer was added to the retina and incubated at 37 °C for 15 minutes to release live cells. Post-incubation, the single-cell solution was centrifuged at 200xg and the papain was deactivated with ovomucoid solution (15 mg ovomucoid (Worthington biochemical) and 15 mg BSA (Thermo Fisher Scientific) in 10 ml of MEM (Thermo Fisher Scientific)). The remaining retinal clumps were triturated in additional ovomucoid solution and filtered through a 20 nm plastic mesh. This step was repeated until the entire retina was dissociated. The collected cells (>90% viability) were stained with Ready probes cell viability imaging kit (blue/red) containing Hoechst 33342 and propidium iodide (R37610, Thermo Fisher Scientific). For capture, retinal cells were diluted to 3E4/ml with 1× PBS (Thermo Fisher Scientific), RNase inhibitor (NEB, 40 KU/ml) and Cell Diluent Buffer (Takara Bio, Cat. #640167)

### The ICELL8™ based single cell capture, single-cell RT-PCR, and library preparation

Using ICELL8 MultiSample NanoDispenser (WaferGen Biosystems), precisely 50 nl (3E4/ml) of the live cell suspension was dispensed into an ICELL8 nanowell microchip containing 5184 wells (150 nl capacity) as described previously (57). Assuming a Poisson distribution frequency for the number of cells per well, about 30% of the nanowells were expected to contain a single cell under optimal conditions. Automated scanning fluorescent microscopy (using an Olympus BX43 fluorescent microscope with a robotic stage) of the microchip was used to select ~1400 wells containing a single live cell (CellSelect software, WaferGen Biosystems). The candidate wells were manually evaluated for debris or clumps as an additional quality control step. The chip was subjected to freeze-thaw to lyse the cells and 50 nl of reverse transcription and amplification solution (following the ICELL8 protocol) was dispensed using the MultiSample NanoDispenser to candidate wells. SCRB-seq was then performed on the candidate wells using a pre-printed primer that adds a well-specific barcode to the (A)n mRNA tail of cellular transcripts. The cDNA libraries were pooled, concentrated (Zymo Clean and Concentrator kit, Zymogen) and purified (using 0.6x AMPure XP beads). A 3’ transcriptome-enriched library was made using the Nextera XT Kit and a 3’-specific P5 primer and sequenced on the Illumina HiSeq 2500 platform. Individual FASTQ files obtained were demultiplexed using the well-specific barcode identifiers for downstream data processing and analysis.

### ScRNA-seq data processing and analysis

#### Mapping and data processing

For single-cell RNA-seq data analysis, FASTQ files were generated from Illumina base call files using bcl2fastq2 conversion software (v2.17). Sequence reads were aligned to the mouse genome mm10 (GRCm38) using STAR alignment software (https://github.com/alexdobin/STAR, version 2.5.1b) with default parameters.

SAM files by STAR were converted into BAM files using SAMtools (http://www.htslib.org). For transcriptome analysis, aligned reads were counted per gene using HTseq (https://htseq.readthedocs.io, version 0.6.0) using default parameters. Next, we used the Seurat package (https://satijalab.org/seurat/, version 2.3.4) for data analysis including quality control, normalization, dimensional reduction, clustering, and visualization. Cells with less than 15% of total expressed genes, and genes that were detected in less than 3% of total cells in each dataset were discarded from all further analyses. In addition, cells with over 20% mitochondrial gene percentage were excluded for filtering of dead or poor-quality cells. After the filtering stage, 1,510 cells and 7,993 genes for the 1 month dataset, 861 cells and 7,858 genes for the 3 month dataset, and 1,224 cells and 9132 genes for the 6 month dataset were used for downstream analysis.

### Normalization and data scaling

After filtering unwanted cells from the dataset, a global-scaling normalization method ‘LogNormalize’ in Seurat was used to normalize the data. In this step, gene expression measurements were normalized by the total expression, multiplied by a scale factor (10,000 by default), and then the data were natural log-transformed.

### Dimensionality Reduction and visualization

Principal component analysis (PCA) was done using variable genes defined as described in the Seurat manual (Butler et al., 2018). Highly variable genes (HVGs) were described by calculating the variance and mean for each gene in the dataset using FindVariableGenes in the Seurat package with default parameters. Genes were sorted by their variance/mean ratio and the top 1000 HVGs were used for PCA. The graph-based method was used to cluster cells. To reduce the high-dimensional gene expression matrix to its most important features, we used PCA to change the dimensionality of the dataset from (cells x genes) to (cells x M) where M is a user-selectable number of principal components. After running PCA, the significant PCs were determined within the first elbow in Jack Straw in datasets. Then, PCA-reduced data were transposed and visualized into 2D space plot using t-SNE, a nonlinear dimensionality reduction method. To cluster cells, the graph-based method was performed using the FindCluster function with resolution 0.8.

### Identifying cluster-specific genes and retinal cell subtypes

We identified cluster-specific genes with the Wilcoxon rank sum test using the FindAllMarkers function of Seurat with min.pct =0.25 and other default parameters, selected the top 20 genes in a specific cluster compared with others at an adjusted p-value < 0.05, and visualized the data with a heatmap. Clusters were assigned by known retinal cell-type markers (Sup. Table 2). In addition, we assessed the expression level of retina-specific markers in clusters using the violin plot. Moreover, we used the FindMarkers function of Seurat to identify DEGs between *Spata7* mutant and wild-type rod cells among the 1, 3, and 6 month datasets by adj p-val < 0.01.

### Functional analysis with Gene Ontology

To functionally characterize the subpopulations, we performed Gene Ontology analysis using significantly DE genes among subgroups of cells. Significantly enriched biological process terms of Gene Ontology were identified using the topGO R package (http://bioconductor.org/packages/release/bioc/html/topGO.html).

### Integration of datasets among different time points

We used Seurat to merge 1, 3, and 6 month datasets to remove batch effects in multiple experiments. First, we identified 1000 HVGs from each dataset using the FindVariableGenes function independently to identify a common source of variation and performed canonical correlation analysis (CCA) with the union of the HVGs from the data sets using the AlignSubspace function of Seurat. Next, with a single integrated data set, we performed dimensional reduction using t-SNE with 10 CCs, clustered cells with the HVGs using the findCluster function (0.6 resolution and 10 dimensions), identified cluster-specific genes using the FindAllMarkers function and assigned retinal cell subtypes using known cell-type markers. After defining retinal cell types we selected the rod cell population and performed a Wilcoxon rank sum test to identify differentially expressed genes (DEGs) among different conditions and time points using the FindMarkers function. DEGs were visualized with heatmap and violin plots with Seurat.

### Single cell trajectory analysis

We used the Monocle 2 package (ver.2.6.1, http://cole-trapnell-lab.github.io/monocle-release/) to generate pseudo-time trajectories for 1, 3, and 6 month old rod cells from *Spata7* mutants and wild-type animals which were extracted from the integrated analysis to discover developmental transitions (Cacchiarelli et al., 2018). We excluded low-quality cells using the detectGenes function of Monocle2 with min_expr =0.1. To normalize expression across cells we applied the estimateSizeFactors function and estimateDispersions function with default parameters. We then performed dimensional reduction using ‘DDRTree’ and the minimum spanning tree was plotted using the plot_cell_trajectory and plot_complex_cell_trajectory functions in Monocle2. To identify DEGs between branches we used the differentialGeneTest function of Monocle2 by adj_p-val < 0.05. Enriched biological process terms of Gene Ontology were identified using the topGO R package.

### Cell cycle analysis of retinal cell populations

To assess whether the clustering assignments were affected by differences in cell cycle phases, we used the CellCycleScoring function of Seurat to predict cell cycle phase based on the expression of G2/M and S phase marker genes (Tirosh et al., 2016). Subpopulation one contains a significantly lower number of cells in the S phase (synthesis) compared to subpopulation two (“Proliferative”; Fisher’s exact test, P < 0.01).

### scATAC-seq data processing

Single Cell ATAC libraries were generated from 7 month old wild-type retinal cells and 6 month old *Spata7* mutant retinal cells. The 10x Chromium Demonstrated Protocol for Nuclei Isolation for Single Cell ATAC Sequencing (Document CG000169) was followed to isolate nuclei. Libraries were constructed with Single Cell ATAC reagent and chip kits following the procedure in the user guides for the Chromium Single Cell ATAC Library & Gel Bead Kit and the Chromium Chip E Single Cell ATAC Kit (Document CG000168). Single cell ATAC library was quantified using a KAPA DNA Quantification kit (KAPABiosystems) and sequenced with 1% PhiX on Illumin Novaseq with the following parameter: read1: 50bp, index1: 8bp, index2: 16bp, read2: 49 bp. After libraries were sequenced, the data from both reagent chemistries were analyzed using the Cell Ranger ATAC pipeline v1.1.0 (https://support.10xgenomics.com/single-cell-atac). We processed scATAC-seq data using the Cell ranger ATAC pipeline. Briefly, this included barcode processing, alignment, duplicate marking, peak calling, cell calling, dimensionality reduction, clustering, t-SNE projection, peak annotation, and differential accessibility analyses. Using the Cell Ranger ATAC pipeline (Figure 5) we performed data aggregation and generated post-normalization unique fragments for wild-type and mutants with 70,546,301 (median=5207/cell) and 102,304,463 (median=5206/cell) UMIs, respectively. Then, open chromatin regions in a genome-wide manner were determined by peak caling using the ZINBA-line mixture model in the Cell Ranger ATAC pipeline (Rashid et al., 2011). Next, a peak-barcode (cell) matrix was generated consisting of the counts of fragment ends within each peak region for each cell for downstream analysis such as dimensional reduction, clustering and visualization. To find differentially accessible motifs between groups of cells, Cell Ranger ATAC uses a Negative Binomial (NB2) generalized linear model to discover the cluster-specific means and their standard deviations and then employs a Wald test for inference. For comparing significant features between mutant and wild-type rods, we used “feature type with Promoter sum filter genes” option in Loup Cell Browser (ver 3.1.0), which are considered features that have an average occurrence greater than one count across the entire data set using the ‘All significant promoter sums per cluster’ option after removing features with a low average count.

### Statistical analyses

Alternative splicing analyses were performed on RNA-seq data from *Spata7* mutant samples and cluster-specific genes in the scRNA-seq profiles were identified with the Wilcoxon rank sum test using FindAllMarkers in Seurat with min.pct =0.25 and adjusted p-value < 0.05. To diminish the false-positive rate, we employed multiple hypothesis correction wherever significance was evaluated across multiple statistical tests. We used at least 5% FDR threshold for significance.

### Data access

The raw and processed data in this study have been submitted to GEO (https://www.ncbi.nlm.nih.gov/geo) with GSE136880.

### Immunofluorescence and histological analyses

Mouse retinae were fixed in freshly prepared Modified Davidson’s Fixative overnight at 4°C (Latendresse et al., 2002). Post-fixation, retinae were processed by a series of ethanol dehydration steps (50%, 70%, 90%, and 100%) and then embedded in Parafilm and sectioned (7 µm; Microtome, Leica) along the vertical meridian. For immunohistochemistry, slides were deparaffinized and antigen retrieval was performed by boiling sections for 30 minutes in 0.01 M Tris, EDTA buffer (pH 9.0) for the detection of GNB3, or 0.01 M citrate buffer (pH 6.0) for detection of histone modifications. Once cooled, slides were washed in PBS and blocked for 1 hour at room temperature in hybridization buffer (10% normal goat serum, 0.1% Triton X-100, PBS). Sections were then incubated overnight at 4°C with corresponding primary antibodies in hybridization buffer. Slides were then washed in PBS with 0.1% Tween and incubated with secondary antibody (Bethyl Laboratories) diluted in hybridization buffer at room temperature for 1 hour. Sections were counterstained with DAPI (1 µg/ml) and mounted using ProLong Gold antifade reagent (Thermo Fisher Scientific) to reduce bleaching. Representative images were acquired on an AxioObserver Z.1 inverted fluorescence microscope with Apotome.2-based optical sectioning and structured illumination on a 100x enhanced chemiluminescence Plan Apochromat objective (Zeiss). Images were processed with ZEN 2 core software (Zeiss). Image stacks were taken with a z distance spanning the entire depth of the cell body for histone modifications in the ONL layer (using DAPI as a reference for each nuclear layer) and projected as a maximum-intensity image.

### Intravitreal injection of TSA

Animals were anesthetized with an intraperitoneal injection of ketamine/xylazine/acepromazine drug combination (37.6 mg/mL ketamine, 1.92 mg/mL xylazine, and 0.38 mg/mL acepromazine) and kept warm during injections. TSA was diluted to 100 nM in DMSO. Intravitreal injections of TSA were performed at P14 in one eye while the other eye was sham injected with an equal volume of DMSO as a contralateral control. Injections were performed with 1 µl of 100 nM TSA to have a final concentration of 20 nM, assuming the free intraocular volume of a mouse eye to be 5 µl (Trifunović et al., 2016). A second dose was administered at P21 and retinae were harvested at P35 for assessment of retinal degeneration by hematoxylin and eosin (H&E) staining.

## Author contributions and acknowledgements

R. D. designed the study, performed the experiments, analyzed the data, and wrote the manuscript. S. K. performed analysis of scRNA-Seq and scATAC-Seq and wrote the manuscript. Y. L. performed scRNA-Seq and scATAC-Seq experiments. S. F. and H. L. performed intraocular injections, R. C. conceived and supervised the research, interpreted the results, and wrote the manuscript. All authors have read and accepted the manuscript. We thank Dr. Graeme Mardon for his valuable insights. This work is supported by grants from Retina Research Foundation (R.C.) and NEI (R01EY018571 and R01EY022356 to R.C.). Bulk and single-cell RNA-Seq were performed at the Single Cell Genomics Core at BCM partially supported by NIH shared instrument grants (S10OD018033, S10OD023469) and P30EY002520 to R.C.

---

**Sup. Fig 1:**
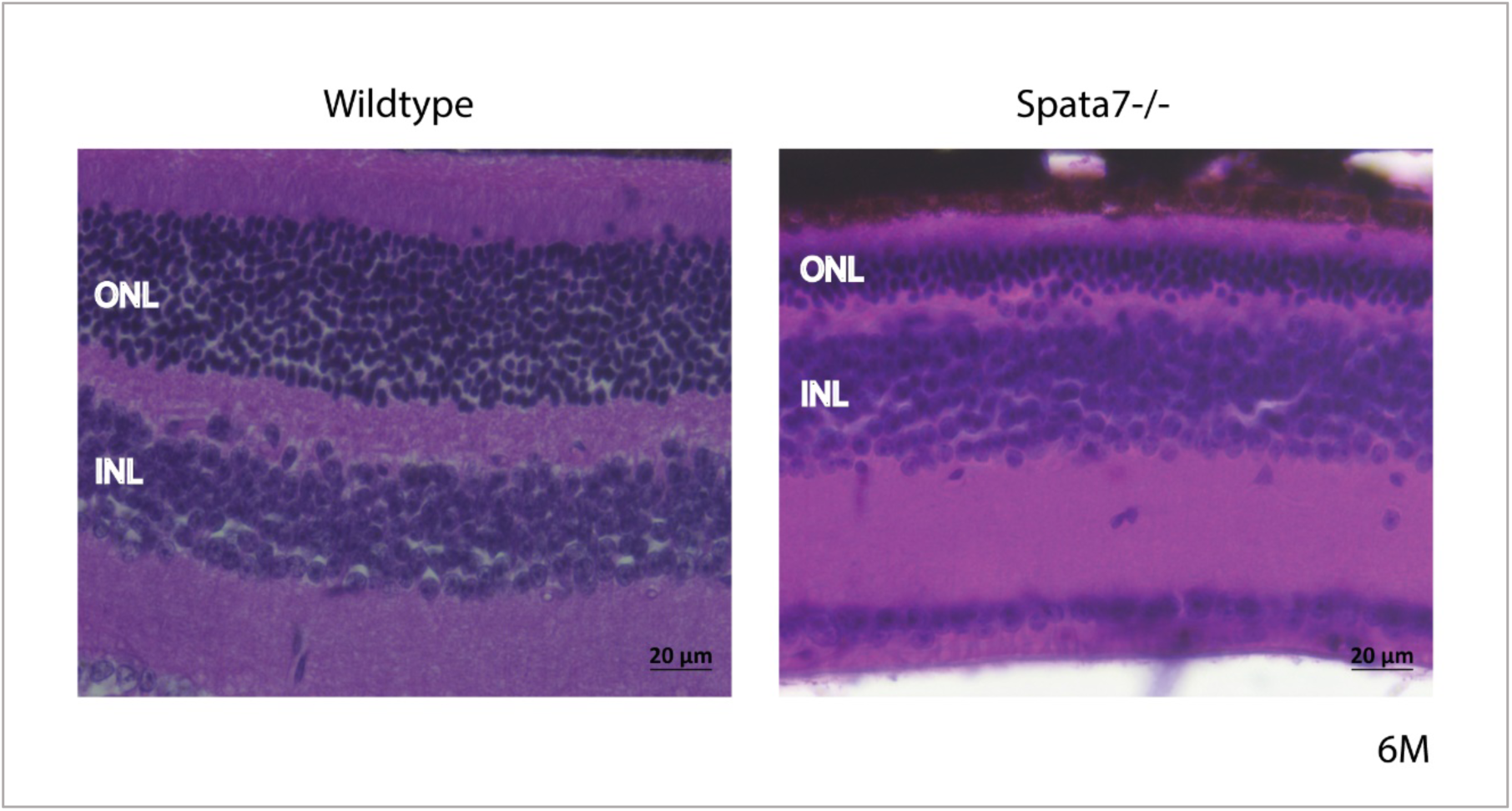
*Spata7* mutant retinas at 6 months still contain many surviving photoreceptors in the outer nuclear layer. H&E of central retinal sections displaying number of surviving photoreceptors in the ONL of the *Spata7* mutant retina at 6 months (6M) of age in comparison to wild-type retina. ONL- outer nuclear layer; INL- inner nuclear layer. Scale bar: 20 µm.

**Sup. Fig 2:**
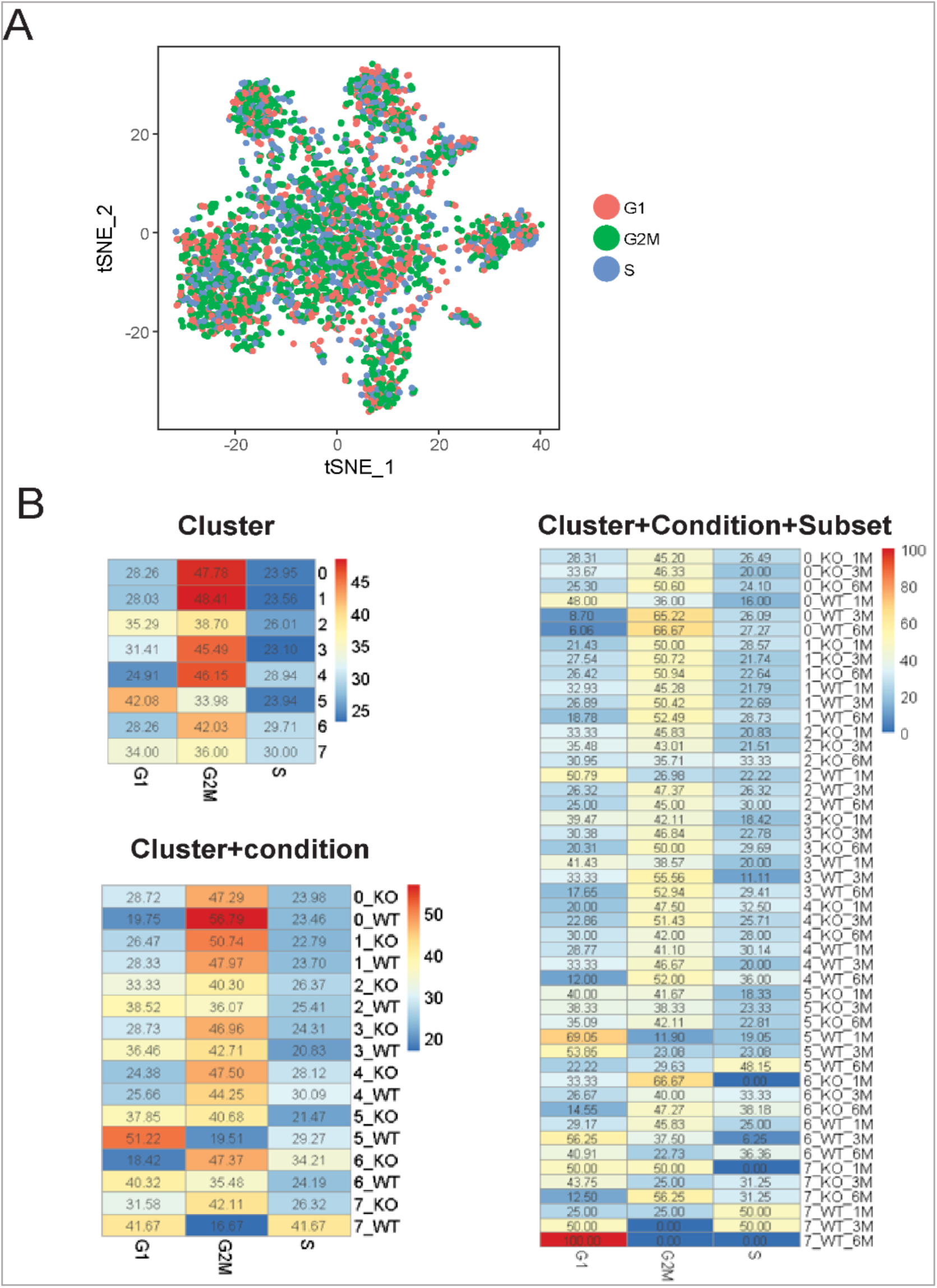
Cell cycle stage prediction indicates high proportion of G2M cells across WT and *Spata7* mutant. Cell cycle stages for single cells in the integrated dataset were predicted by subpopulation using the CellCycleScoring function of the Seurat package. A. Cell cycle stages were visualized on the t-SNE plot. B. The proportions for cells assigned in each subgroup were calculated based on subgroups cell cluster, genetic condition, subset categories, and generated heatmaps.

**Sup. Fig 3:**
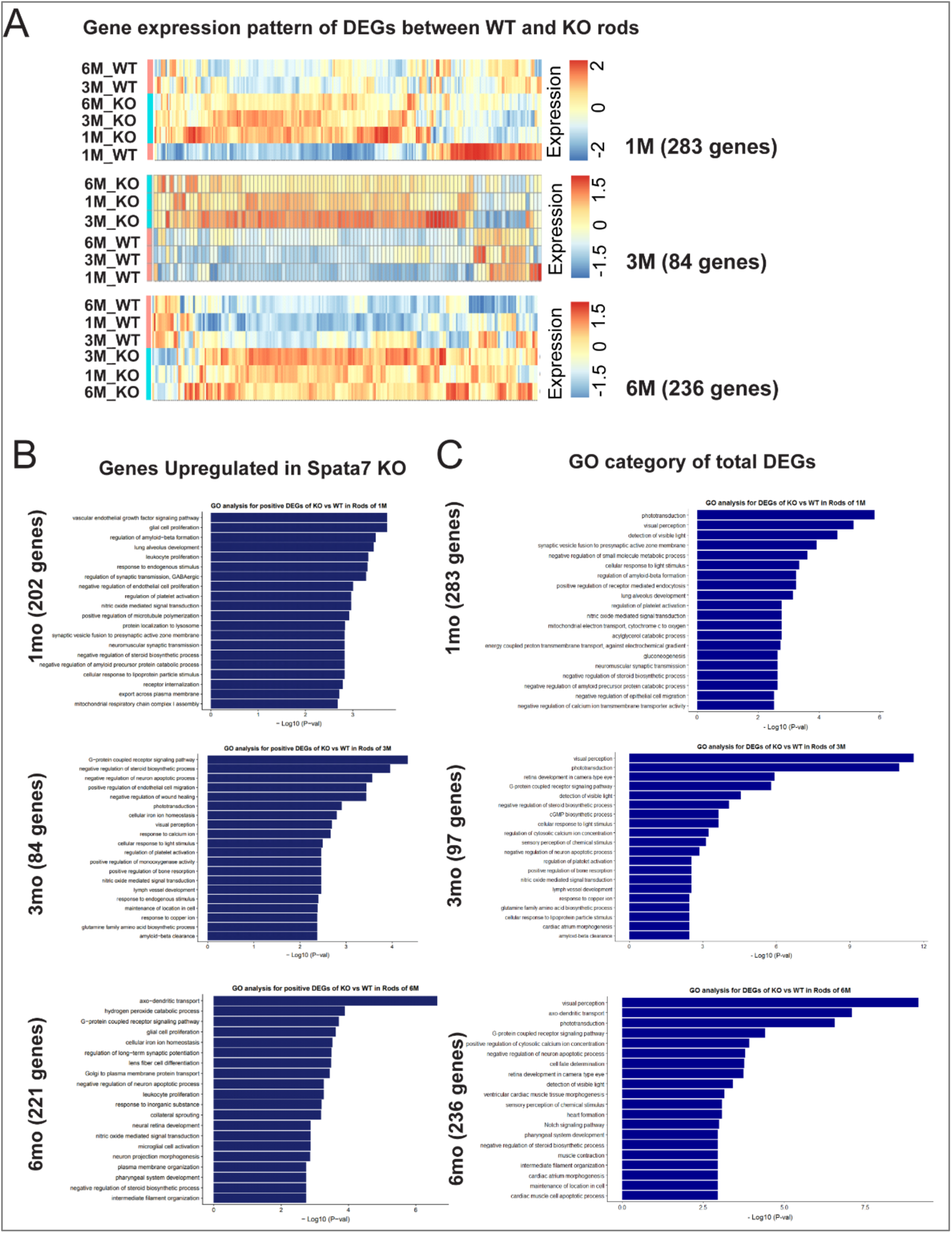
DEGs between sub-clusters of WT and *Spata7 mutant* cells. A. DEGs between subclusters were identified using the differentialGeneTest function of Monocle2 with FDR < 0.01. B. Gene ontology category for genes upregulated in *Spata7* mutants C. Gene ontology category for all DEGs

**Sup. Fig. 4:**
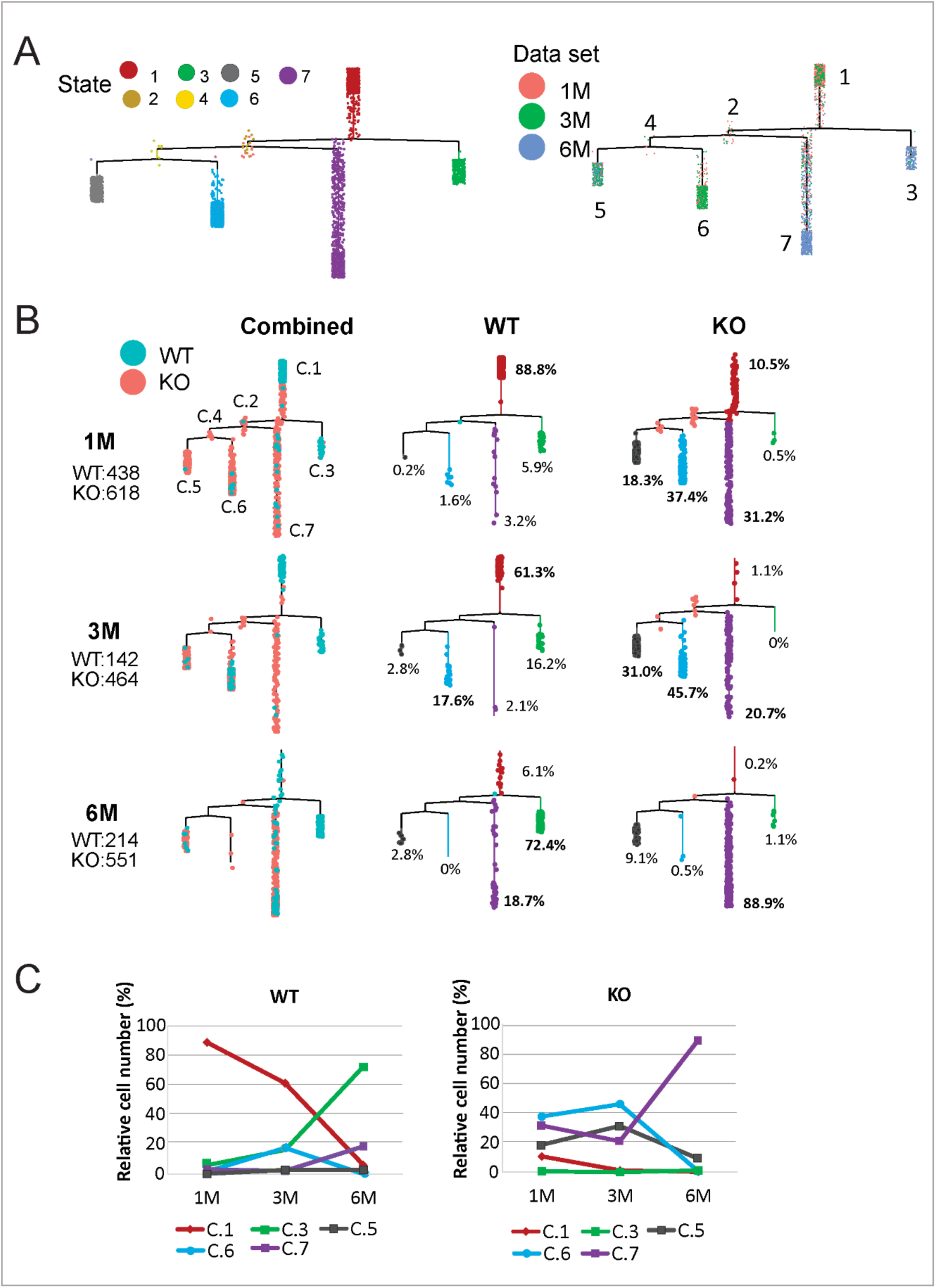
Different lineages are observed in *Spata7* mutant rods during degeneration. A. Trajectory analysis for rod cells. B. Monocle2 recovers a branched single-cell trajectory beginning with rod cells and terminating at different time points. Each dot represents a cell. The color describes the genetic background in the first panel and the branch identity in the second and third panels for *Spata7* mutant and wild-type, respectively. C. Proportion of *Spata7* mutant and wild-type rod cells in each branch in 1, 3, and 6M datasets.

**Sup. Fig 5:**
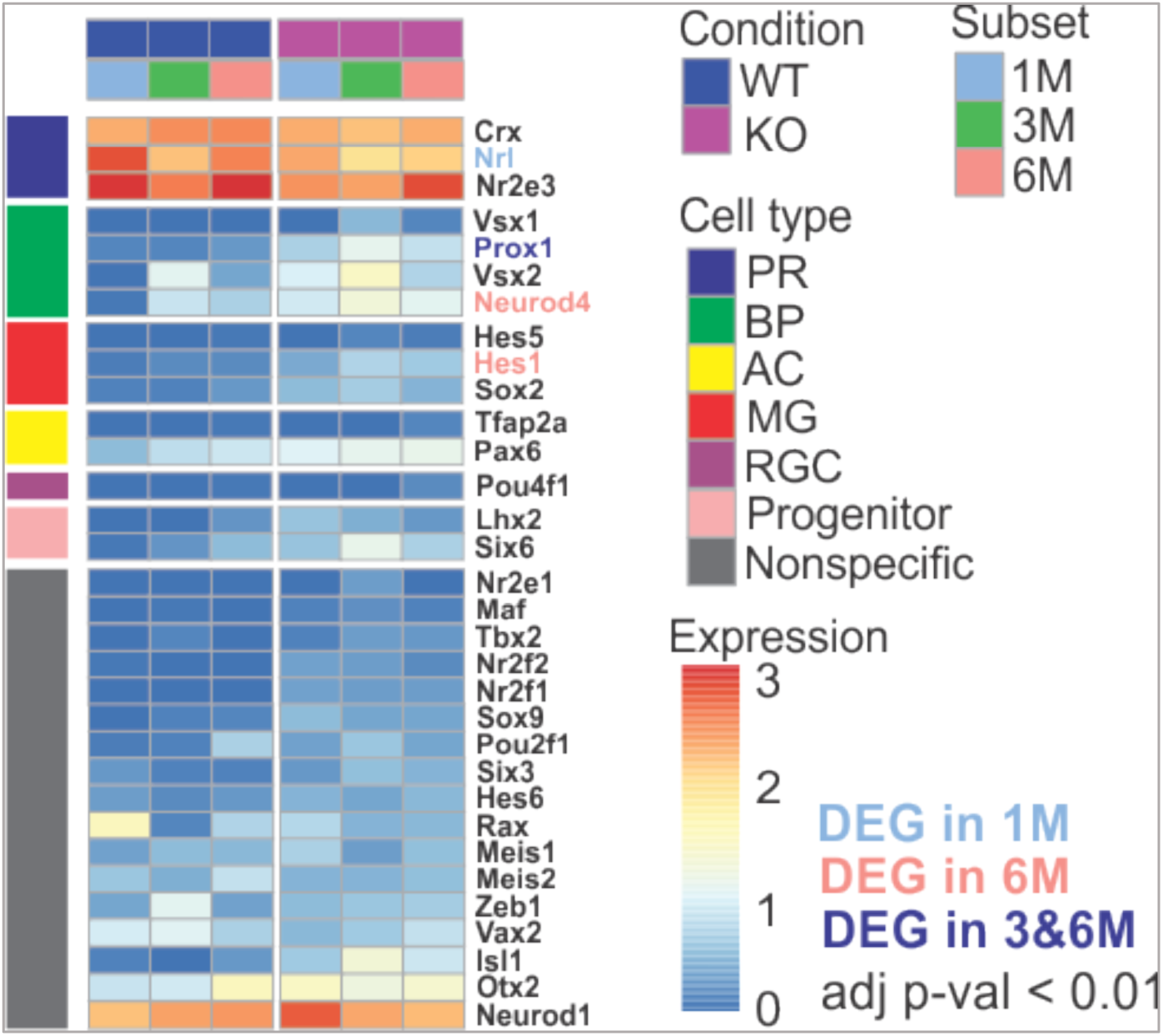
Expression levels of major retinal transcription factors across the dataset remain largely unchanged. Heatmap describing the average expression of genes of sub-groups by condition and subset in transcription factor genes was visualized with heatmap. Expression values are median centered after log2 transformation.

**Sup. Fig 6:**
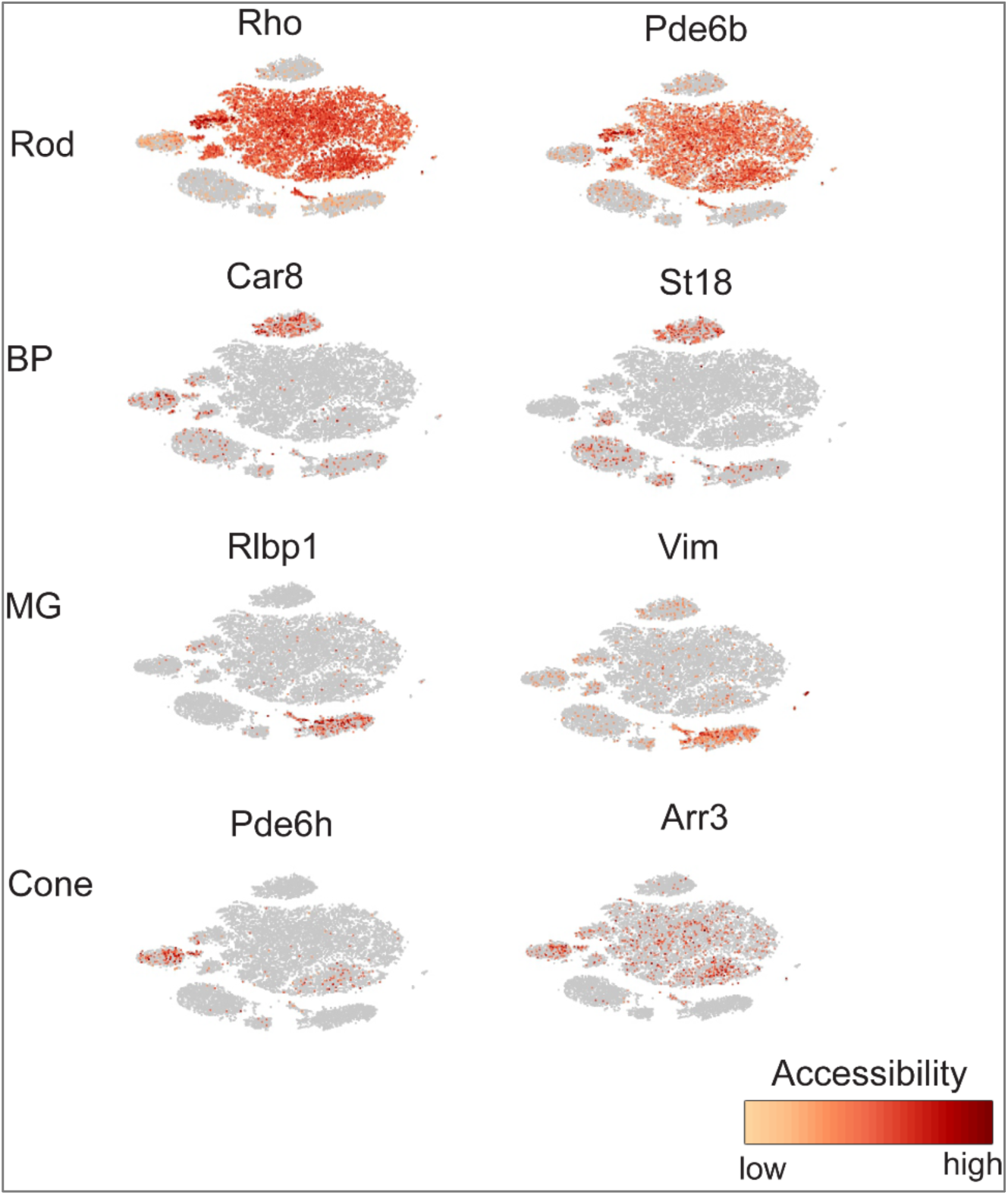
Chromatin accessibility of known marker genes on combined scATACseq data. To assign clusters to specific retinal cell types, known marker genes such as *Rho* and *Pde6b* for rod, *Car8* and *St18* for BP, *Rlbp1* and *Vim* for MG, and *Pde6h* and *Arr3* for cone were used on Loupe Cell Browser (https://support.10xgenomics.com/single-cell-atac/software). MG; Müller glial cell, and BP; bipolar cell.

